# Robust, long-term video EEG monitoring in a porcine model of post-traumatic epilepsy

**DOI:** 10.1101/2022.01.17.476340

**Authors:** Luis Martinez-Ramirez, Andrea Slate, George Price, Ann-Christine Duhaime, Kevin Staley, Beth A. Costine-Bartell

## Abstract

To date, post-traumatic epilepsy (PTE) research in large animal models has been limited. Recent advances in neocortical microscopy have made possible new insights into neocortical PTE. However, it is very difficult to engender convincing neocortical PTE in rodents. Thus, large animal models that develop neocortical PTE may provide useful insights that also can be more comparable to human patients. Because gyrencephalic species have prolonged latent periods, long-term video EEG recording is required. Here, we report a fully subcutaneous EEG implant with synchronized video in freely ambulatory swine for up to 14 months during epileptogenesis following bilateral cortical impact injuries or sham surgery The advantages of this system include the availability of a commercially available system that is simple to install, a low failure rate after surgery for EEG implantation, radiotelemetry that enables continuous monitoring of freely ambulating animals, excellent synchronization to video to EEG, and a robust signal to noise ratio. The disadvantages of this system in this species and age are the accretion of skull bone which entirely embedded a subset of skull screws and EEG electrodes, and the inability to rearrange the EEG electrode array. These disadvantages may be overcome by splicing a subdural electrode strip to the electrode leads so that skull growth is less likely to interfere with long-term signal capture and by placing two implants for a more extensive montage. This commercially available system in this bilateral cortical impact swine model may be useful to a wide range of investigators studying epileptogenesis in PTE.

**Significance:** Post-traumatic epilepsy (PTE) is a cause of significant morbidity after traumatic brain injury (TBI) and is often drug-resistant. Robust, informative animal models would greatly facilitate PTE research. Ideally, this biofidelic model of PTE would utilize a species that approximates human brain anatomy, brain size, glial populations, and inflammatory pathways. An ideal model would also incorporate feasible methods for long-term video EEG recording required to quantify seizure activity. Here, we describe the first model of PTE in swine and describe a method for robust long-term video EEG monitoring for up to 14 months post-TBI. The relatively easy “out-of-the-box” radiotelemetry system and surgical techniques described here will be adaptable by a wide array of investigators studying the pathogenesis and treatment of PTE.

## Introduction

Post-traumatic epilepsy (PTE) is the most common form of acquired epilepsy, which is often refractory to treatment. PTE is usually defined as two seizures at least 2 weeks after the traumatic brain injury (TBI) event. In humans, the latent period prior to the development of epilepsy can last several months or longer (Annegers, Hauser et al. 1998). The mechanisms of epileptogenesis during this latent period are unknown. Models of PTE have been mainly restricted to variations of rodent impact models where an impact is made to the thin cortex overlying the hippocampus (impact, fluid percussion injury, weight drop) (Pitkänen and McIntosh 2006, Ostergard, Sweet et al. 2016, Keith and Huang 2019) with very limited work in gyrencephalic species (Friedenberg, Butler et al. 2012, Steinmetz, Tipold et al. 2013).

The majority of human PTE is neocortical in origin (Gupta, Sayed et al. 2014). Although processes that are widely hypothesized to be epileptogenic occur in rodent neocortex after trauma (Jin, Huguenard et al. 2011), to date it has not been possible to develop a robust neocortical PTE model in rodents (Bolkvadze and Pitkänen 2012, Pitkänen, Lukasiuk et al. 2015). This may be due to the small size of the rodent brain, which results in significant epileptogenic hippocampal injury when the neocortex is damaged by trauma (Komoltsev, Sinkin et al. 2020), or it may reflect a relatively high threshold for the development of stable, chronic epilepsy in the rodent neocortex (Chang, Yang et al. 2004, Cela, McFarlan et al. 2019).

The interspecies differences in brain anatomy may also underlie the signficant species differences in responses to therapy. In TBI, over 150 therapies have been shown to reduce lesion volume in rodent models but have failed to demonstrate efficacy in humans or in swine (Margulies and Hicks 2009). Differences in brain anatomy including location of the hippocampus (direct mechanical trauma in rodent impact models vs. secondary cascades or diffuse TBI in humans), variability in the abundance of white matter and white matter injury, variation in the pathoanatomic character of the lesion developing from the injury (cavity in rodents vs. a remodeled area with thick gliotic scarring in gyrencephalic species), the population and characteristics of the glia (Azevedo, Carvalho et al. 2009, Herculano-Houzel 2009, Khakh and Deneen 2019, Khrameeva, Kurochkin et al. 2020), differences in the matrisome (Pokhilko, Brezzo et al. 2021), the degree of genetic variation among individual subjects of a given species, and immune response differences (Seok, Warren et al. 2013, Warren, Tompkins et al. 2015) all are host factors that may affect the development of PTE among species, and thus may affect our understanding of the development of PTE in humans. Indeed, brain size and the duration of the latent period are positively correlated (Lillis, Wang et al. 2015). In order to design therapeutic interventions that may prevent PTE in humans, the constellation of injuries that occur in humans must be modeled in a brain more similar to humans. Models utilizing gyrencephalic species may bridge this gap.

Simplified ex vivo models of PTE such as slice preparations are also available (Berdichevsky et al. 2012, 2016; Goldberg and Coulter 2013). Because of the severity of injury (complete transection of the hippocampus at 350 um intervals), 100% of explants develop medically intractable PTE (Berdichevsky et al. 2016). These preparations are very amenable to longitutinal microscopy studies (Lillis et al. 2015; Lau et al. 2021) but they are based on rodent hippocampi, not neocortex. Further, the complete penetrance of epilepsy complicates studies of epileptogenic mechanisms.

A technical barrier in the adoption of large animal PTE models is the need for reliable, long-term EEG monitoring because of the longer latent period compared to rodents (Lillis, Wang et al. 2015). Long-term monitoring of large animal models of epileptogenesis has been limited to date.

Non-human primates have been used for long-term EEG monitoring but are expensive (Vuong, Garrett et al. 2020), and the use of livestock species for research is more acceptable to the general public perception than non-human primates or companion animals. Here, we describe extreme long-term monitoring of a swine model of PTE using a video EEG radiotelemetry system. We discuss the advantages and disadvantages of a contusion model in swine monitored with a commercially available video EEG radiotelemetry system enabling real-time analysis.

## Materials and Methods

### The Radiotelemetry System

The Data Sciences International PhysioTel Digital radiotelemetry system enabled transmission of EEG data from subcutaneous implanted electrodes in real-time via Bluetooth to a transceiver connected to a communication link controller (CLC) that managed the EEG implants and relayed digitized data to a computer. Video was recorded and synchronized to the EEG data. At the peak of the study, up to 6 swine were recorded at the same time.

All EEG and video acquisition hardware was installed via the manufacturer’s instructions (DSI Implantable Telemetry System Manual (International 2017) (**Figure 1**). The communication link controller (CLC) allows communication between the implants and computer by discovering implants, assigning frequencies, and relaying digitized EEG data via an ethernet connection to the computer. The CLC can manage up to 6 EEG implants at a time. The CLC was housed in the animal room in a stainless-steel lock box to prevent water damage during cage cleaning. The CLC was connected via ethernet cables tunneled through the ceiling to the data acquisition computer located just outside the animal room. The EEG implant communicated to the CLC via transceivers. Per bank of pens, 2-3 transceivers were secured to the animal cage walls at a height that was inaccessible to the animals while also not blocking the field of view of the video cameras. The transceiver cables were routed along the outer perimeter of the cages and along the room walls to the CLC. It is recommended to place transceivers at right angles to one another to minimize areas of poor signal reception and prevent signal drop-off (International 2017). Three video cameras (AXIS M1145-L Network Cameras, Axis Communications, Lund, Sweden) were installed on the ceiling 2 - 3 feet away from the bank of pens and placed in a way that maximized field of view at each of the three banks (**Figure 2**). Each camera recorded two independently housed or three socially housed animals at a time. Each camera was enclosed within an acrylic box (a modified basketball display case, 10-1/4” sq. x 10-1/4” h, The Container Store; not provided by DSI) to protect the cameras from water damage during pen washing. It was opened at night to allow video recording in infrared mode. All cables from the video cameras were routed along the ceiling to an ethernet data port in the animal room to an ethernet data port outside the animal room and connected to the data acquisition computer. Feeders fixed to the front of the cage were removed and replaced with rubber dishes on the pen floor to increase visibility with video recording. Video was recorded using Noldus Media Recorder 4.0 software (Wageningen, the Netherlands). A key synchronization step was required to synchronize the video and EEG acquisition software using the “Network Time Protocol” resulting in a 5 ms specificity between the two.

**Figure 1.**
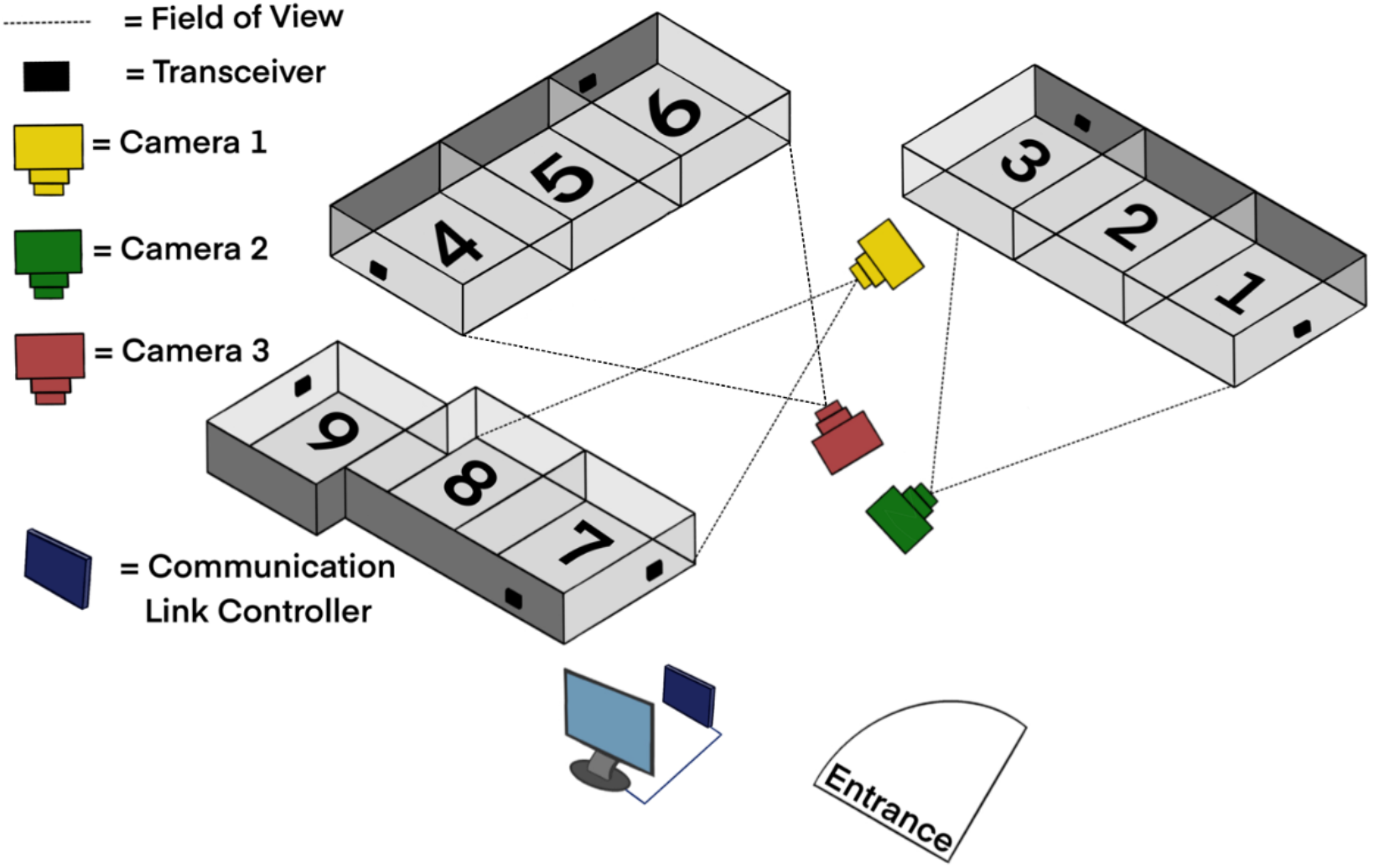
Diagram of the video EEG digital radiotelemetry monitoring unit. The transceivers receive EEG data from the implants via Bluetooth, which are connected via cables (not shown in diagram) to the communication link controller, which then communicates via cable to the computer that stores the data outside of the telemetry room. With 3 cameras, up to 6 pigs could be recorded with video at the same time with large pigs taking two pens in a total of 8 pens. Pigs were separated for their weekly night of video EEG recording.

**Figure 2.**
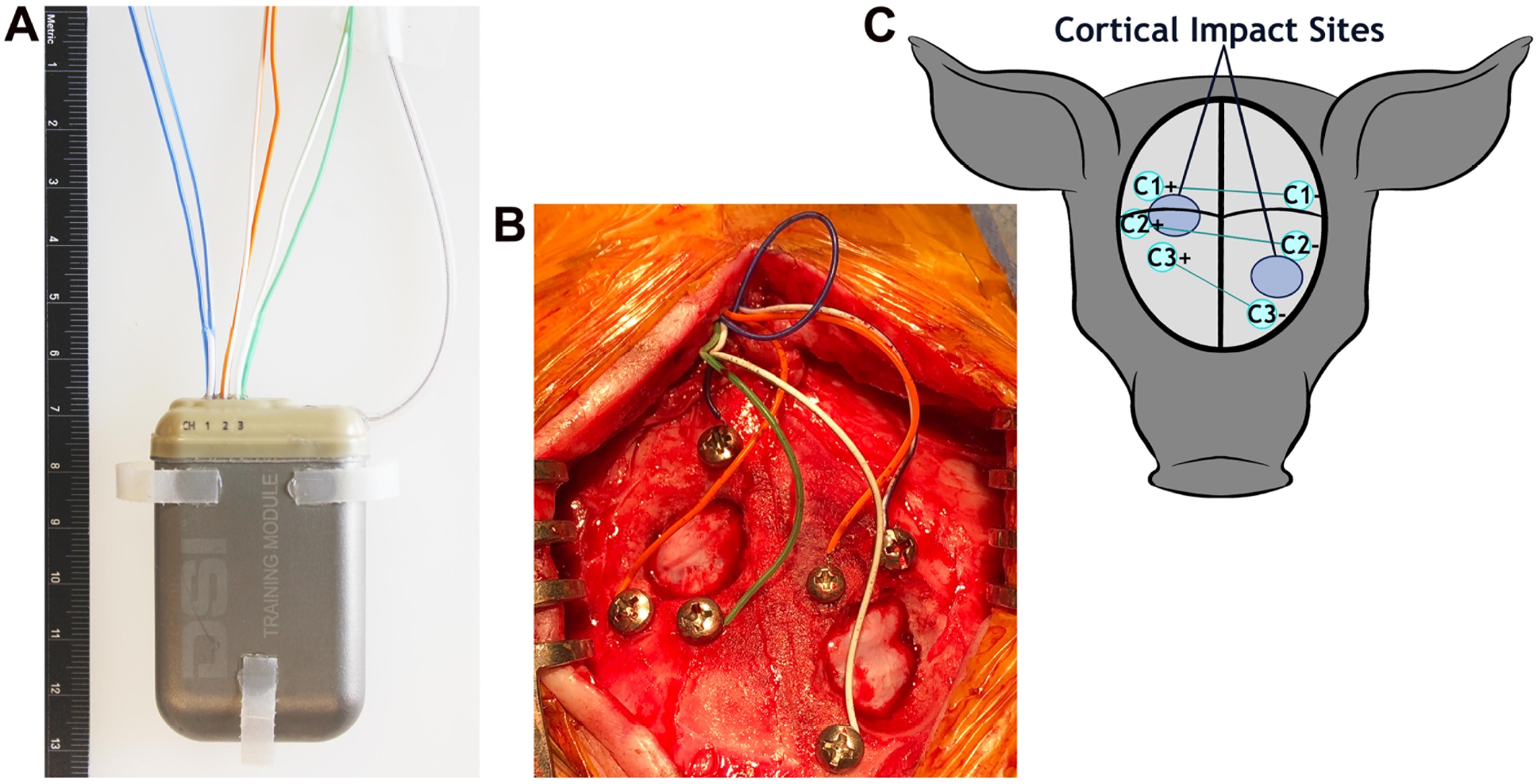
EEG implant and montage. **A.** The 59 x 38 x 15 mm EEG implant with electrodes exiting the top of the implant, the antennae projecting off the right of the implant, tabs on the sides and bottom that are used to suture the implant in place. **B.** Pigs received bilateral cortical impact through the burr holes. The electrode array prior to the application of dental cement. **C.** A schematic of the 3-channel bipolar montage with electrode sites centered around the sites of cortical impact. Black lines represent the sagittal and coronal sutures.

The PhysioTel Digital L03 series implant (3 channels, **Figure 2A**) was used in this study in conjunction with the Ponemah Data Acquisition software (Data Sciences International; DSI, St. Paul, MN). The acquisition frequency was 500 Hz. The L03 series implant had six biopotential leads with no common leads. Instead, each pair of the biopotential leads was coupled into an instrumentation amplifier resulting in differential inputs to 3 channels. Each biopotential channel had a common mode rejection ratio of −40 dB or better at test frequencies of 0.5 Hz and 10 Hz. The common mode signal applied to the biopotential channel was generated with respect to the implant housing connection (ground). There were no hardware filters. A software filter that was a 30^th^ order finite impulse response was applied. A low pass filter for 150 Hz was used to guarantee accurate signal acquisition and channel bandwidth of 0.5 Hz - 100 Hz; this was verified by testing with a signal or function generator. No low frequency signal was filtered. A Blackman window was used for calculating coefficients for the filter designed as a windowed-sinc filter (Data Sciences International, personal communication).

In addition to the biopotential lead signals, the implant also provided temperature measurements as well as activity measurements measured via a three-axis accelerometer. “The three-axis accelerometer provides acceleration data along the x-, y-, and z-axes, relative to the orientation of the implant. Acceleration for the x, y and z axes was reported as a value from an analog-to-digital converter. A range of at least −7 Gs to +7 Gs was provided, with a corresponding output from approximately 0 to 4095. A value near 2047 was displayed when zero acceleration for a given axis was sensed when in a steady, neutral alignment (orthogonal) to earth’s gravitational field. The displayed sampling rate for the x, y and z axis acceleration data was 10 Hz. Along with the values from each axis of the accelerometer, Ponemah also reports an activity value calculated from the accelerometer axes…” (Data Sciences International, 2017). The accelerometer data was used to detect movement to screen for convulsions compressing the length of video required for manual screening.

The manufacturer’s estimate for battery life for the implant was 90 days. In order to enable prolonged monitoring, in order to conserve battery life, the implant was turned on and off using a strong magnet swiped over the implant and could also be turned off via the acquisition software.

Animals were housed in a temperature-controlled animal facility with 12-h light/dark cycles. Video EEG was recorded following a schedule that maximized vivarium recording capacity. Animals were warehoused at an off-site facility as space for additional space. When on-site, the animals were recorded with EEG and video or video-only at least every other week and as space permitted. Animal facility staff were given a recording schedule and moved animals per the schedule.

### Surgery for EEG implantation and cortical impact

Male, castrated Yucatan minipigs (Sinclair Bio Resources LLC, produced in Windham, ME; N = 17) were implanted at age 4.92 ± 0.16 months (21.5 ± 0.79 kg; mean ± SEM). Grain pellets were removed the evening prior to surgery and the swine were fasted for 12 hrs. Water was available at all times. The animals received a Hibclens (Norcross, Georgia) bath the day before surgery and the morning of surgery prior to anesthetic induction where the animals were gently sprayed with warm water and scrubbed with approximately 5 ml of Hibclens into a lather. Diazepam was administered (2-4 mg/kg, PO) in simple syrup 30-45 minutes prior to anesthetic induction. The swine were sedated with a pre-anesthetic mix consisting of ketamine (20 mg/kg, IM), xylazine (2 mg/kg, IM), and atropine (0.03 mg/kg, IM). Pigs greater than 50 kg were sedated with Telazol (2.2-4.4 mg/kg, IM), xylazine (2 mg/kg, IM), and atropine (0.03 mg/kg, IM) to minimize total injection volume.

The pig was transported to the operating room under 3-5% isoflurane and oxygen delivered via nose-cone mask while monitoring oxygen saturation and heart rate using a handheld pulse oximeter unit. Before entering the operating room, the swine was shaved at incision sites, feet wrapped with Vetrap (3M Healthcare, Saint Paul, MN), eye lubricant was placed along the inner edge of the eyelid then closed with a piece of tape followed by a full Tegaderm patch (3M Healthcare, Saint Paul, MN). Xeroform petrolatum gauze (Covidien, Mansfield, MA) was inserted into the ears. The swine’s head, ears, and neck were washed with 2% chlorhexidine cloths then moved onto the operating table.

An intravenous (IV) catheter was placed in an ear vein. Vancomycin (10-20 mg/kg) was infused via IV over 30-60 minutes followed by saline (2-4 mL/kg/hr; IV). Swine were intubated and mechanically ventilated with isoflurane titrated to 1-2% mixed with medical air. Ventilation was adjusted so that end-tidal CO_2_ was maintained between 35 and 45 mmHg with a peak pressure of 20-25 mmHg. Core body temperature was measured via a rectal probe and maintained at 37-39°C using a heating pad and Bair hugger forced air blanket. Swine received an infusion of saline (2-4 mL / kg / hour). Blood pressure as measured via a cuff on a hindlimb was maintained above 45 mmHg. Saline boluses (2-4 mL/kg, IV) were administered for hypotension (mean arterial pressure (MAP) < 45 mmHg). Epinephrine (5 ug/kg, IV) was administered if saline did not successfully increase MAP. End-tidal CO_2_, oxygen saturation, blood pressure, heart rate, and core body temperature were monitored and recorded every 15 minutes. Pre-injury and 2-hour post injury blood was collected via IV or superior vena cava venipuncture for later analysis. Blood was collected 24-hour post-injury via the superior vena cava. Buprenorphine (0.02 mg/kg) was administered IM 15 minutes before the first incision.

The ears were wrapped with sterile Vetrap. Tegaderm was placed over the top of the snout, over the eyes, and around the ears creating a continuous perimeter of Tegaderm around the surgical site. The swine was positioned in sternal recumbency such that the head and neck were accessible, rolls of absorbent pads were placed underneath the swine to reduce pressure on the abdomen. The surgery was performed in a single scrub position. The scrub was performed with a separate scrub pack after the surgeon scrubbed and gowned. The incision sites (head and right side of neck) were prepped using surgical sterile technique using 70% ethanol followed by betadine using gauze held with a dedicated scrub hemostat, then with ChloraPrep which was allowed to dry (Becton Dickinson, Franklin Lakes, NJ).

After prepping, sterile Steri-drapes (3M Healthcare, Saint Paul, MN) were placed around the incision sites followed by Ioban (3M Healthcare, Saint Paul, MN). Lastly, a large Tiburon split-sheet sterile drape (Cardinal Health, Dublin, OH) was placed over the entire surgical area. The drape was clipped to an IV stand in front of the animal’s head such that the endotracheal tube remained in view for adjustment when necessary and to test mucous membrane and jaw laxity for anesthetic plane.

To minimize risk of infection with implants, all instruments were autoclaved and gas sterilized instruments were not used. Bupivacaine (1.5-2.5 mg/kg) was administered subcutaneously at the head incision site. The first skin incision was made along the sagittal midline from above the snout to the crown of the head. The skin was detached from the periosteum to expose the skull. The sagittal and coronal sutures were identified and a Hudson drill was used to make a burr hole on the right coronal suture over the rostral gyrus. A dural separator was used to detach the dura from the underside of the skull. Bone ronguers were then used to expand the burr hole to approximately 2 cm in diameter. Hemostasis was obtained with sterile bone wax.

The cortical impactor guide was secured to the skull at each burr hole (Duhaime, Margulies et al. 2000). The cortical impact device was screwed into the guide until the 1.07 cm in diameter tip was just touching the surface of the dura. The indenter was then deployed (over 4 ms) over the closed dura with an indentation velocity of 1.7 m/s (Duhaime, Margulies et al. 2000). In a similar manner, a second burr hole was made on the left, rostral to the coronal suture to expose the very rostral portion of the brain. The prefrontal cortex is the somatosensory cortex that represents areas of the face and portions of the mouth; the rostral gyrus is the somatosensory cortex of the snout (Craner and Ray 1991, Missios, Harris et al. 2009). The offset of contusion locations on the two sides was chosen to minimize any functional disability that might be caused by bilaterally symmetric lesions.

A Stille bone hand drill (Sklar Surgical, West Chester, PA) with 1.5 mm and 2.0 mm drill bits was used to drill six holes for skull screws, three on each side around the area of the burr hole (**Figure 1B**). Everbilt Pan Head Philips Stainless Steel #4 screws (Home Depot Product Authority, Atlanta, GA; not provided by DSI) with a 2.85 mm head diameter were filed down to varying lengths between 5-15 mm to accommodate variable skull thickness. Screws were threaded into the drilled holes with a surgical screwdriver. A dural separator was used during installation of the screws to verify that each screw was placed through the skull and in contact with the dura.

Bupivacaine (1.5-2.5 mg/kg) was administered subcutaneously at the neck incision site on the right side of the neck approximately 6 cm posterior to the ears and 6 cm lateral to the midline. An approximately 5 cm neck incision was made with a different set of sterile instruments that had been set aside and not previously used. A pocket was open under the skin via blunt dissection beneath the subcutaneous fat or under the trapezius muscle until it was slightly larger than the implant. The sterile EEG implant was removed from the sterile packaging using a new set of sterile gloves. The edges of the skin were draped with a second set of Steri-drapes and gauze packed around the edge of the incision such that the implant did not touch the skin. The EEG implant was implanted either under the subcutaneous fat or under the trapezius muscle and sutured into place through the implant tabs. The EEG implant biopotential leads were tunneled under the skin from the implant site to the caudal end of the head incision using a Nelson 35 French trocar (Sklar Surgical, West Chester, PA). The neck incision was irrigated copiously with sterile saline, and the incision was closed with interrupted 2-0 PDS suture (Ethicon, Somerville, NJ) in the subcutaneous layer, and 3-0 Monocryl (Ethicon) subcuticular suture followed by LiquiVet Rapid tissue adhesive (Oasis Medical, Mettawa, IL).

The biopotential leads were trimmed to a size long enough to reach the skull screws while leaving enough additional length to allow for pig growth and movement. Leads were placed in a bipolar montage to the right and left focused around the cortical impact sites (**Figure 2 B,C**). Approximately 5 mm of the silicon insulation was stripped from the biopotential leads and the exposed lead was wrapped around the shaft of each screw and secured using silk suture. The screw was then tightened to secure the lead to the skull. Screws and leads were required to be low profile to allow skin closure. Maxcem Elite dental acrylic (Kerr Corp., Brea, CA) was applied to completely cover the exposed screws and leads to ensure electrical isolation from surrounding tissues (International 2012). Once the dental cement set, the cortex was irrigated with sterile saline and the incision was closed with interrupted 2-0 PDS suture for the subcutaneous layer and 3-0 Monocryl running subcuticular skin closure followed by skin adhesive.

To optimize the time interval over which epileptogenesis could be observed, a subset of pigs who received cortical impact did not receive an EEG implant until 6.43 ± 0.64 months post-cortical impact (27.22 ± 2.59 kg) though they were video recorded. The intended time to implant was 4 months post-cortical impact but was delayed due to lock-out from our facilities due to the COVID pandemic.

Sham pigs (N = 3) underwent the same surgical procedure, including installation of the EEG implant and skull screws etc., except the cortical impact device was not deployed. The scrub and surgery required 6 hours with an additional hour required for recovery.

### Post-surgical recovery and monitoring

Isoflurane was reduced and the pig encouraged to breath by allowing end tidal CO_2_ to increase. Buprenorphine was administered (0.025 mg/kg, IM) and fentanyl transdermal patches (1-4 ug/kg/hr) were placed on the lower back for pain management for 72 hours after surgery. The animal was then transferred to the animal facility under 1-2% isoflurane and extubated. The animal was monitored until ambulatory. Antibiotic ointment (2% Mupirocin ointment; Taro Pharmaceuticals, Hawthorne, NY) was applied to both skin incisions the first day after surgery to prevent infections. Prophylactic cephalexin (10-20 mg/kg) was administered orally three times a day for seven days post-operatively. The animals received twice daily evaluations for three days post-operatively and five times a week until the incisions were fully healed. Subjects were not transferred to a satellite facility until full healing was achieved approximately 1-month post-surgery.

Though the areas receiving cortical impact were expected to be clinically silent, swine were often somnolent the day after surgery, sometimes had temporary difficulty with coordination/movement of the front left leg that resolved in the day or two after surgery. Though no formal consistent vision testing was performed, temporary limitations in vision were suspected as some swine receiving bilateral cortical impact had absent menace responses, startled to touch, and tripped over their food bowl. This behavior was not observed in sham pigs. The signs of vision impairment resolved by post-surgical day 1 or 2.

The site of implantation in the neck displayed significant tissue swelling in the first 3-5 days post-surgery but resolved thereafter. Less swelling was observed when the implant was placed under the trapezius muscle vs. under the subcutaneous fat.

### Animal Husbandry Procedures

In these long-term experiments, the animal enrichment team provided swine with regular stimulation and socialization. Staff provided a new toy or activity daily. Once swine reached 50 kg, they were placed in two pens to enable space to accommodate their larger size. Swine were introduced to one another over time so that compatible swine were socially housed during days of video-only recordings. The specific individuals in each 24-hour video were logged in a spreadsheet where they were identified by physical markings or the presence of two ear tags vs. one. When pigs were scheduled to have EEG and video recorded, they were placed in an individual pen. Isolating the pig during EEG recording was crucial in later analysis as it was difficult to assess if an event was real electrographic activity or artifact due to their pen partner’s movements. Metal feeders in front of the cage were removed and replaced with rubber feeders during healing from surgery and during recording as they obstructed the view from the video cameras. Placards were posted in the room instructing staff where to scratch the pigs to avoid interfering with the incision sites. Swine received regular food treats including yogurt and apricots and often did not require restraint for pre-anesthetics due to acclimation with our study staff.

Animals that received a second ear tag for identification purposes for recording with social housing or required hoof trimming (approximately every 3 months) were anesthetized for these procedures. Swine were removed from feed 12 hours before anesthesia and were anesthetized using the pre-anesthetic mix described above (or, once greater than 50 kg, swine were given Telazol; 2.2-4.4 mg/kg, IM) and then anesthetized under 3-5% isoflurane and oxygen. Hair was clipped if necessary and was cleaned with alcohol or betadine. The tags were cleaned with alcohol or betadine and inserted into the outer portion of the ear while avoiding the outer cartilage supporting the ear and the central ear vein and other large veins using an ear tag applicator and/or the hoofs were trimmed. After the procedure was completed, isoflurane was reduced, and the pig was encouraged to breathe and recovered.

Yucatan skin required regular management. To treat dry, itchy skin, staff applied mineral oil to the pig’s body daily until their skin was healthy, and thereafter, weekly to maintain healthy skin. Regular oiling prevented the animals from scratching their implant site with their hindlegs or against the side of the cage. Several Yucatan spontaneously developed blisters and open lesions over the dorsum consistent with bullous pemphigoid as previously described in this strain (Mirsky, Singleton et al. 2000, Olivry, Mirsky et al. 2000). The open lesions were cleaned with diluted chlorhexidine gluconate and treated with topical antibiotics as needed. Many displayed allergic reactions to bacitracin, neomycin, and polymyxin topical antibiotics as well as to unidentified substances during surgeries. Allergic reactions were limited to the skin, were self-limited, and did not require treatment.

### Euthanasia and brain collection

After developing PTE or at 12-14 months post cortical impact (76-86 kg), pigs were NPO for 12 hours, given diazepam (2-4 mg/kg, PO), Telazol was administered (2.2-4.4 mg/kg, IM) 30 minutes later then deeply anesthetized with 3-5% isoflurane and intubated with a 9-10 endotracheal tube. The pig was moved by 4 staff members on a pig board, transferred to a hydraulic lift, motorized cart, and moved to a down draft necropsy table. After ensuring a surgical plane of anesthesia, swine were euthanized via exsanguination by transcardiac perfusion with 0.9% saline and 10% formalin. The skull was opened with a bone saw and the brain, including the olfactory bulbs and 2 cm of spinal cord were collected. The time required for euthanasia, carcass disposal, clean-up, and brain removal was 8 hours. The brain was weighed and post-fixed at 4°C for 5-7 days. The cerebral hemispheres were coronally sliced. Blocks were paraffin embedded and stored in sealed containers at room temperature for future investigation.

## Results

### Electroencephalographic Recordings

To date, this is the first published model of any type of epilepsy in swine and provides the longest-term recording of which we are aware. The system was relatively easy to set up.

Video EEG was recorded until PTE developed or for 12 months (maximum of 14 months) in 13 subjects: 10 injured and 3 sham pigs. The average duration from time of EEG implantation to the end of the experiment among all animals was 11.5 months achieving very long-term monitoring.

A portion of the pigs receiving cortical impact developed epileptiform spikes, electrographic seizures and convulsions. The rate of epilepsy and analysis of interictal epileptiform discharges in relation to onset of convulsions will be published in a separate manuscript. The EEG of injured pigs who developed PTE showed a wide array of epileptiform discharges similar to those seen in human patients. Simple and complex spikes, sharp waves, spike trains, clusters of waves and spikes, and electrographic seizures were recorded in multiple injured animals prior to and after developing convulsions (**Figure 3**).

**Figure 3.**
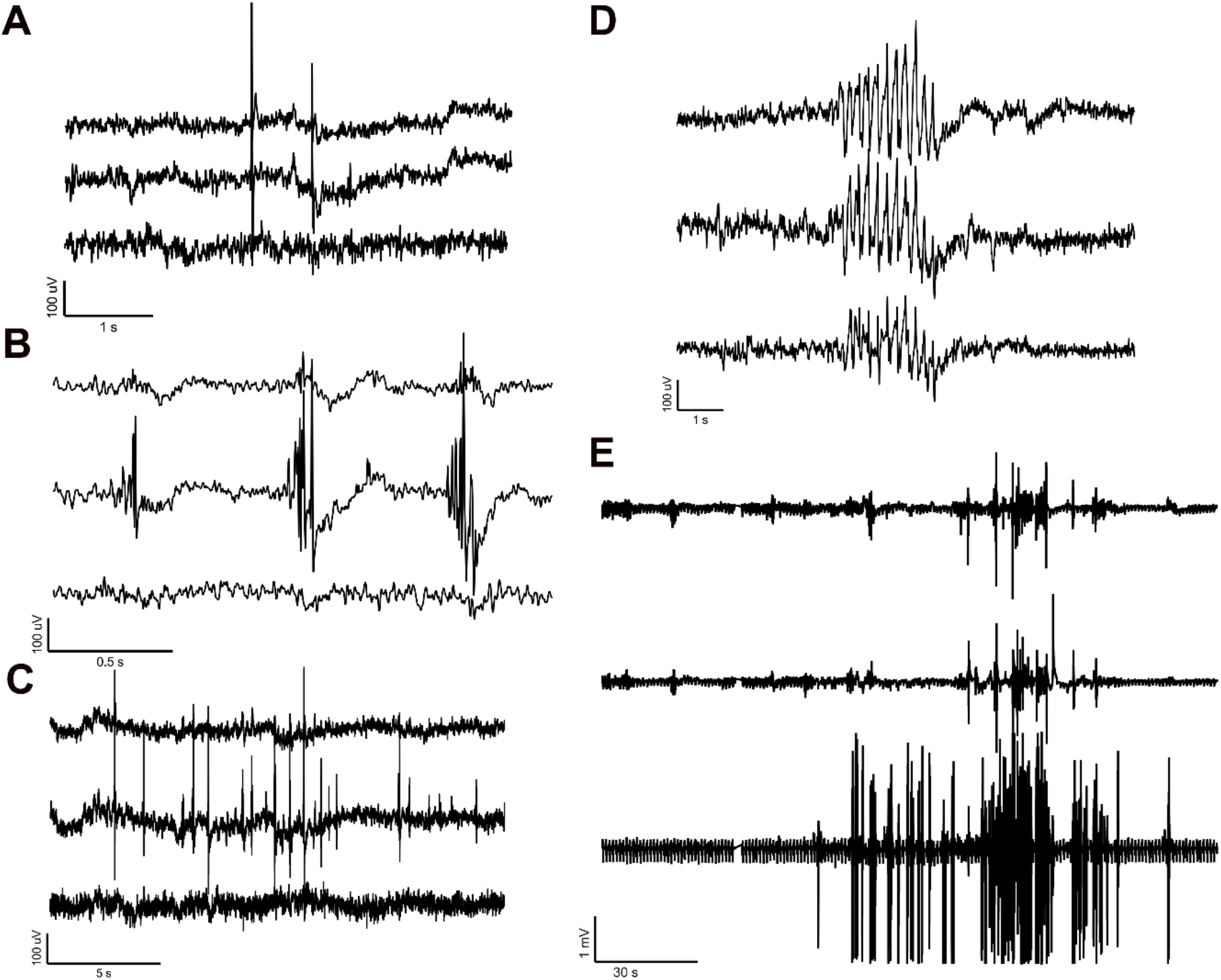
Interictal spikes and electrographic seizure on our 3-channel array. **A**. A simple spike. **B**. Complex spikes. **C**. A train of complex spikes. **D**. Clusters of waves with spikes. **E**. An electrographic seizure accompanied by tonic-clonic convulsions.

Similar to ambulatory EEG systems in rodents and humans, movement artifact occurred during large movements verified by the synchronized video (**Figure 4**), but the system was robust and did not display movement artifact from minimal muscle movement. Large amplitude and high frequency sharp spikes saturated the EEG signal when staff fed the swine. Large amplitude artifact occurred when animals jumped up on the sides of the pen to greet neighboring animals or facility staff, playing with enrichment toys or scratching floorboard, and during headshakes. Low amplitude muscle artifact occurred with eating. However, most low speed activities such as drinking water, moving the head around during normal voluntary movement, walking around the pen, and gentle sleep-rocking were not detected on the EEG. As a result of these factors, periods where the animals were lying down or doing minimal physical activity were optimal for EEG analysis.

**Figure 4.**
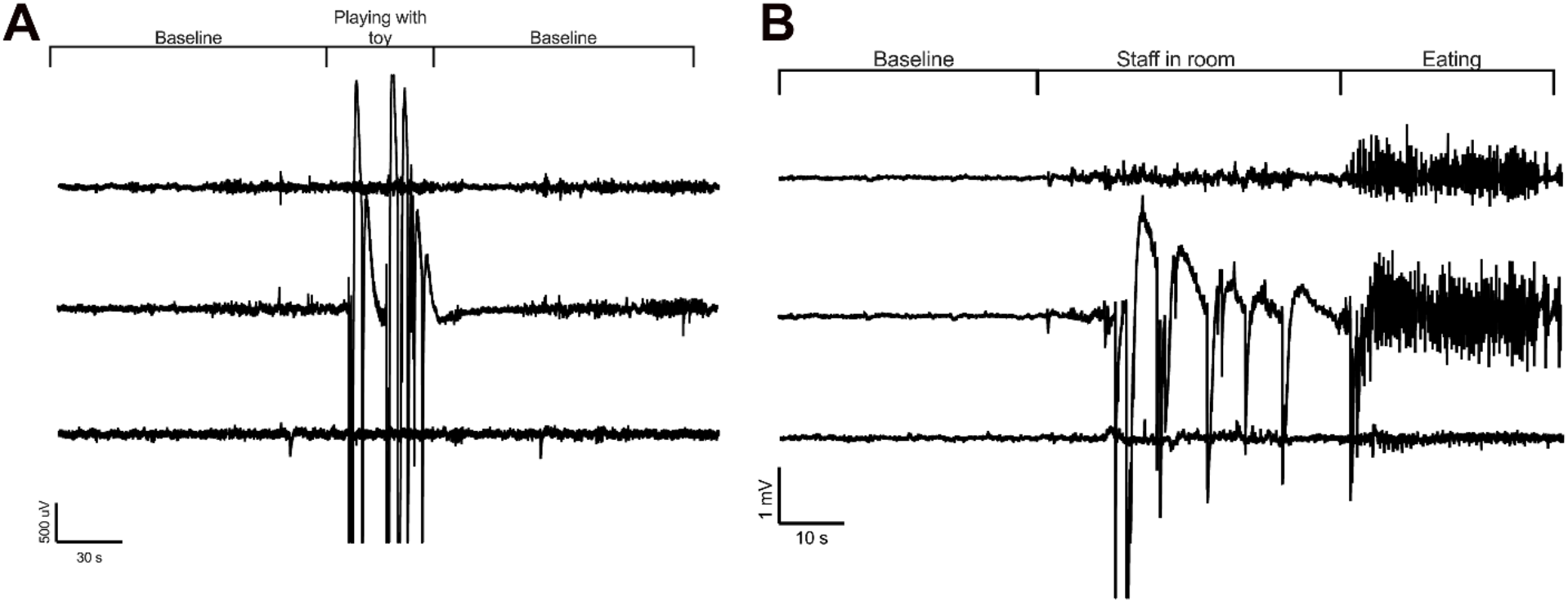
Artifact on our 3-channel array. **A**. High amplitude artifact was observed while the animal was playing with a toy or during a vigorous head shake (not shown). **B**. High amplitude artifact in response to anticipation of feeding (including jumping up on side of pen) followed by low amplitude muscle artifact of eating.

### Behaviors

Throughout the study, injured pigs were observed to have a variety of stereotypical peri-ictal behavioral repertoires. These behaviors included staring spells, head nodding and shaking, instances of forelimb clonus and tonic extensions, lip smacking and licking, and tonic-clonic convulsions followed by post-ictal stillness. Some of the epileptic animals seemed aware of the oncoming epileptic episodes such that they would gently lower themselves to the ground or lean against the side of the cage to lower before convulsing. Most pigs would convulse during emergence from sleep. In only one instance was a pig observed to drop suddenly. The analysis of semiology for each pig with PTE as well as non-peri-ictal behavior specific to pigs with PTE vs. normal swine behavior will be published separately. Unlike rodents with Racine scale of level 5, no individual was observed to be apneic, none reared up on its back legs, nor died from convulsions. In collaboration with our large animal veterinarians, no convulsive event was deemed to endanger animal welfare. The only injury observed was abrasions on their side and legs from rubbing on the floor during convulsions.

### EEG implant limitations

A limitation of this telemetry unit is the limited battery life of 90 days, which limits recording to 1-2 times/week in subjects in which it may require several months to develop PTE. Three implants indicated several days of battery life left but failed to record at the very end of battery life; actual battery life was typically 80 days when switched on and off regularly. One of 13 EEG devices completely stopped functioning 20 weeks into the study. Despite having approximately 78 days of battery life remaining, the implant failed to turn on. The subject was changed to a video-only recording schedule. An attempt was made to use the manufacturer’s crimping tool to switch out an implant where the battery was expended and splice the existing biopotential leads to a fresh new device. However, this surgical procedure was not possible as wires were encapsulated by the body and difficult to remove without damaging. The animal was switched over to a video-only recording schedule.

### Limitations in Large Animal Housing

Another limitation of this study were the logistical issues of housing these large animals at our institution for up to 14 months. A great deal of planning and communication with the animal facility was needed to successfully accommodate these animals. However, given that the animal facility had limited space, many of our study animals had to be sent out to an outside animal facility which resulted in week-to month-long gaps in video EEG recording for some animals. Due to collaboration with our animal housing facility administration, a contract with an outside institution was established so that future studies will have the ability to record video EEG at the warehouse site to ensure consistent capture of data.

Space restrictions resulting in limitations in consistency of video EEG recording for large animals staying for prolonged periods of time could be overcome by installing an additional DSI system at a warehouse facility where staff can manage recording as described here as well as exporting data. Swine could be sent to the warehouse recording facility when healed from the surgery (3-4 weeks post-surgery). In this scenario, up to 12 swine could be recorded weekly.

Staff compliance with the recording schedule was high, while compliance with removing metal feeders during video EEG recording was low. Additionally, compliance with keeping the transponder so it was not located on the pen between the pig and the video camera was also low. An auditing schedule could improve compliance and improve the quality of the video acquired.

### Montage Limitations

Though we were able to quantify epileptiform activity, it is not possible to re-montage the array limiting our ability to identify a specific seizure focus with 3 biopotential channels. The acquisition bandwidth was limited to 0.5 Hz to 100 Hz thereby limiting the ability to acquire infraslow frequencies or high-frequency oscillations (< 0.5 Hz, > 100 Hz respectively).

There was significant accretion of skull bone over time that resulted in unpredictable shifts in screw placement and dura contact from implantation to the end of the study. At age 17 months, the nuchal ridge is very large over the crown of the head and paranasal sinuses cover the skull on the sides the skull covering the rostral portions of the brain. Screws placed most caudally were most affected by skull thickening often becoming completely embedded in skull bone (**Figure 5A**) while rostral screws sometimes were found to be exposed to air in the sinuses and were no longer contact with the dura. Some screws were found to leave shallow indentations in the brain transversing the dura in some cases (**Figure 5B**) but were not associated with the incidence of PTE: both pigs with or without PTE exhibited screw indentations. Despite the alterations in screw location, we were able to record long-term from the pigs in all but one subject. In one subject, the ability to detect spikes was absent at approximately 34 weeks postinjury due to the screws becoming completely embedded into the skull.

**Figure 5.**
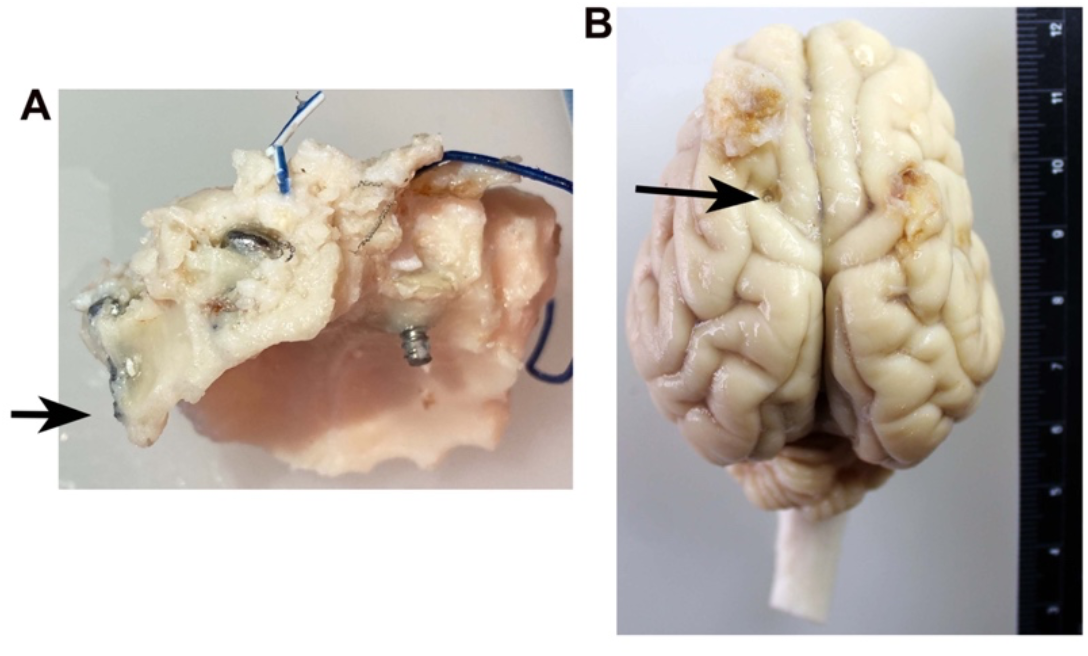
The evolution of screw location. Because of the significant accretion of skull over time with the development of the nuchal ridge, screws initially placed to just contact the dura eventually were completed embedded within the skull (**A**; arrow), exposed to air in the sinuses, or embedded within the brain (**B;** arrow). Screw indentations were observed in both swine that did and did not develop PTE and did not appear to be a cause of PTE.

### Morbidity and Mortality

Early in the project, we encountered problems with infections that were solved by adapting surgical sterile technique protocols from the human operating room along with other strategies (**Table 1**). The EEG implant is large and provides an extensive area ripe for development of biofilm. Implementation of the protocols in Table 1 and regular pig oiling completely ended the infection issue. As many measures were implemented at once, it is impossible to identify which were the key factors, but once the infection problem had abated, one pig that failed to receive its pre-surgical baths developed a MRSA infection. Therefore, pre-surgical baths may be key to preventing infections. Specifically, 4 pigs were infected at the implant site and required early euthanasia and were excluded from the study. The infection in two pigs was due to Methicillin-resistant Staphylococcus aureus (MRSA) infections that developed rapidly (2 weeks) after implantation. Two pigs had infections that presented later at the implant site at 7 and at 14 weeks after implantation though post-surgical swelling was greater than normal. These infections were positive for *Beta-hemolytic Streptococcus, Staphylococcus schleiferi*, and *Streptococcus porcinis*. In one instance, the delayed infection may have resulted from a superficial scratch early after implantation that eventually abscessed. The scratches may have resulted from the pig scratching from dry, irritated skin. Thereafter, swine were on a regular oiling schedule to prevent itch and thus scratching.

**Table 1.**
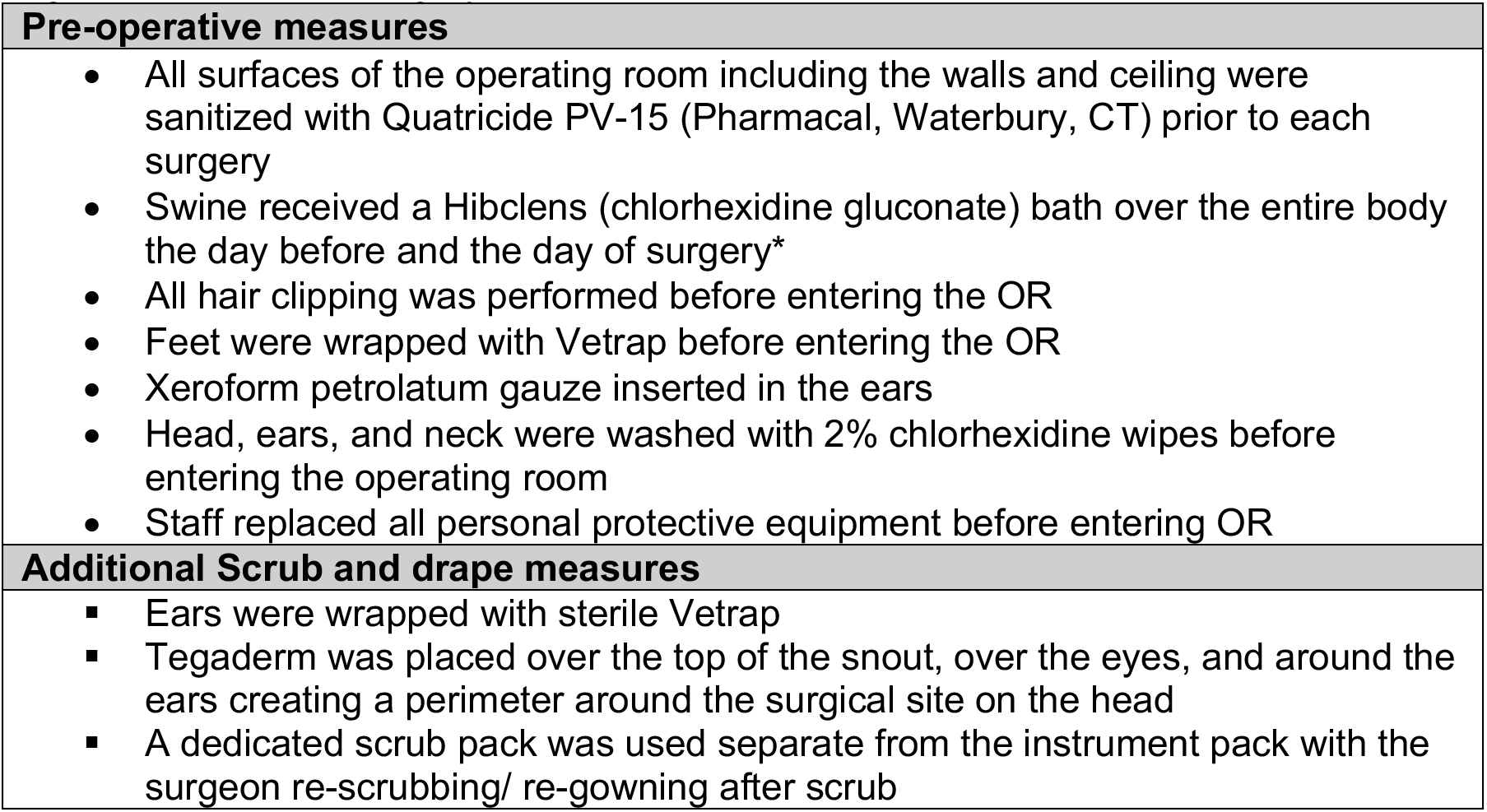
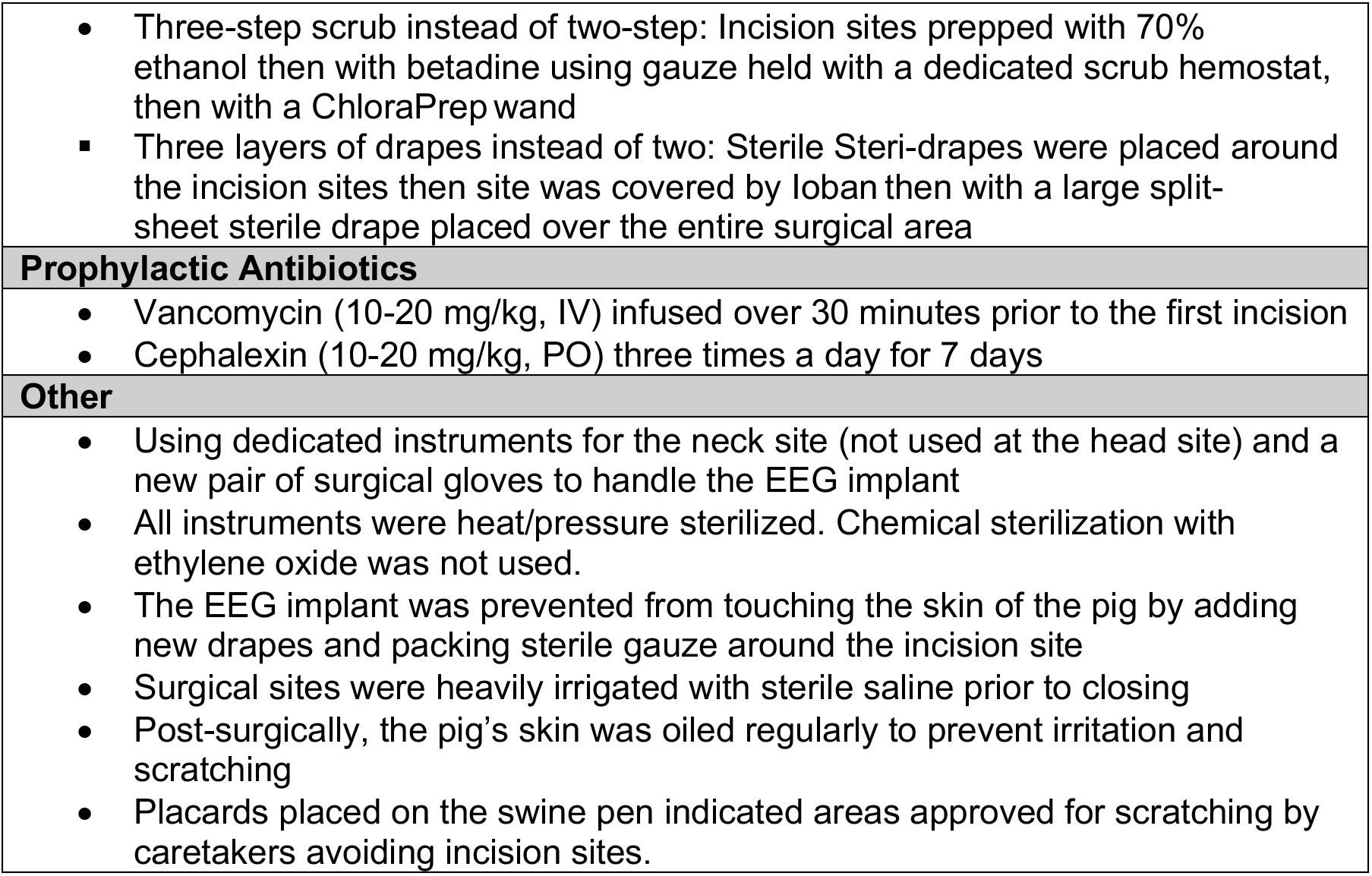
Procedures and medications initiated to prevent infection above and beyond standard large-animal survival surgery standards.

## Discussion

Acute EEG recordings in anesthetized swine after various brain insults is relatively simple and is well-established (Costine-Bartell, Price et al. 2021), but recording EEG in awake and restrained and/or tethered swine over the course of several days is more difficult. The standard of EEG in PTE in rodent models of TBI is tethered EEG (Shandra and Robel 2020) or radiotelemetric EEG (White, Williams et al. 2010). In rodents, the onset of PTE is 21 days (or earlier) to around 2 months post-TBI. However, in large-brain species, epileptogenesis after TBI develops over several months, requiring a corresponding period of video EEG monitoring and analysis (Lillis, Wang et al. 2015). Piglets can be recorded with scalp electrodes for EEG or amplitude-integrated EEG while restrained and kept calm for durations of time similar to clinical EEG in humans. Such scalp recordings enable the study of acute effects of TBI or therapies for brain insults (Wang, Zhang et al. 2014, Atlan and Margulies 2019, Barata, Cabañas et al. 2019). Similarly, telemetric EEG devices can be temporarily placed on the head allowing the piglet to ambulate freely while awake or sleeping (de Camp, Dietze et al. 2018). However, the labor required to apply, record, and remove EEG may be cost prohibitive if extended to several months, and may also interfere with normal behavior. The first report of recording from chronically implanted EEG electrodes in swine was up to 3 months (Stromberg, Kitchell et al. 1962). Depth electrodes into the hippocampus have been used up to 6 months in tethered swine with head mounts (Forslid, Andersson et al. 1986, Ulyanova, Cottone et al. 2019). Recent advances have been made in telemetry in swine with a DSI implant to record EEG in fully ambulatory EEG in piglets for up to 5 days (Rault, Truong et al. 2019). In this instance, EEG was limited to a single channel but was sufficient for power analysis (Rault, Truong et al. 2019). Such short-term recordings do not require strict and extensive peri-operative and operative sterile implantation techniques for success (Rault, Truong et al. 2019).

While electrodes are generally resistant to infection (Stromberg, Kitchell et al. 1962), fully unrestricted ambulatory EEG requires an implanted transmitter and battery assembly that comprises a large-volume foreign body that is prone to infection seeded at the time of implantation. The advantage of the fully subcutaneous neck implant is the lack of risk of infection after initial implantation. In contrast, head mounts have a continuous risk of infection due to repeated butting and consequent wound dehiscence. Exteriorized head mounted systems are preferable in non-human primates as they pick at their subcutaneous implants (Vuong, Garrett et al. 2020) and do not typically head butt. However, swine head butt frequently and thus frequently break exteriorized head mounted systems, resulting in a constant risk of infection and/or destruction over time. Here we report measures that were successful in preventing infection at implantation and report long-term stability of the fully subcutaneous implant for 12-14 months allowing for extreme long-term monitoring.

We observed a loss of signal at 34 weeks in one subject and implant malfunction at 20 weeks in another. However, the duration of recording exceeded what has been reported for hippocampal depth electrodes tested in naïve swine, where there is a significant loss of oscillation power within the first month followed by persistent loss over 6 months (Ulyanova, Cottone et al. 2019). Additionally, hippocampal depth electrodes might create more damage than the alterations inflicted by skull screws as they are associated with acute hemorrhage and chronic lesions with activated microglia and gliosis (Ulyanova, Cottone et al. 2019). Many types of EEG arrays may perturb the system that they record, and certainly, the integrity of the signal is an issue in all methods of invasive, long-term EEG recording.

The advantages of this system are 1) availability of a “kit” where DSI provides most of the equipment, 2) excellent signal-to-noise ratio with algorithms that create reference and ground resulting in minimal movement artifact, 3) the swine freely ambulate and the implant is completely under the skin/not at risk of destruction, and 4) availability of continuous video with excellent synchronization of video to EEG. The disadvantages of this system are: 1) the skull screws may become embedded in the skull or exposed into the air of sinuses as the skull undergoes significant accretion and remodeling, resulting in the loss of channels over time, 2) the screws may become embedded in the brain, which occurred in both swine that developed PTE and those that did not develop PTE, and 3) the system does not allow re-montage of the electrode array. Export of the EEG to other universal data file formats is necessary for advanced analysis.

Potential alterations to this system to address limitation disadvantages 1) and 2) could include using EEG subdural electrode strips spliced to the DSI implant using the DSI splice kit. These electrodes would slip underneath the dura. Disadvantage 3) could be addressed by installing one telemetry implant in each side of the neck/hemisphere allowing a montage with additional electrodes. With these improvements, skull screws would be avoided post hoc re-montaging and advanced signal analysis of the recording would be possible.

## Conclusions

The methods described in this study demonstrate the feasibility of using swine to model post-traumatic epilepsy via video EEG using a commercially available radiotelemetry system. This system allowed for up to 14 months of monitoring producing good quality EEG. The set up was largely uncomplicated and required minimal upkeep of successfully implanted animals. This robust system may be of benefit to detect epilepsy in swine over the long period of epileptogenesis in this species. Slight modifications to this system as described may overcome the significant skull accretion in swine and improve the quality of EEG acquired.

## Author Contributions

KS and BCB designed the experiments, LMR, AS, GP, ACD, BCB performed the experiments, LMR, BCB, and KS analyzed the data, LMR, BCB, and KS wrote the manuscript.

## Acknowledgements

The authors would like to thank Jason Zhao, Andrew Ding, John Dempsey, Akhila Penumarthy, Tess Del Prado, Bohan Sun, Nancy Yang, Anna Akkara, and Ashley Buggy who assisted with deep cleaning the OR, surgeries, post-surgical assessments, and perfusions. We thank Tawny Stinson for the illustration. We are sincerely grateful for the assistance of Dr. Donna Jarrell and Titilayo Lamidi for navigating the space issues and working with us to find solutions. We are especially thankful for the help of Amanda Zall and Kimberly Degrenier and the staff at the Center for Comparative Medicine for all the tasks going above and beyond regularly animal husbandry including: their assistance with moving pigs to the correct pens, removing feeding bins, covering and uncovering the video cameras, pig baths, pig oiling, pig enrichment and socialization, making sure our equipment appeared to be functioning, and observations of the animals’ well-being. We thank Troy Velie at DSI for answering our technical questions.

